# Relocating the congocidine biosynthetic cluster to the central chromosomal compartment enhances gene expression and antibiotic production in its native host, *Streptomyces ambofaciens*

**DOI:** 10.64898/2026.03.04.709497

**Authors:** Nicolas Delhaye, Hoda Jaffal, Thomas B. Grégory, Hervé Leh, Jean-Luc-Pernodet, Sylvie Lautru, Stéphanie G. Bury-Moné

## Abstract

*Streptomyces* bacteria are renowned for their intricate life cycle and prolific production of specialized metabolites, including antibiotics. Their linear chromosome is spatially compartmentalized: the central region contains highly conserved and expressed genes, while the terminal regions harbor less conserved, poorly expressed sequences, often rich in specialized metabolite biosynthetic gene clusters. To investigate the relationship between genome architecture and gene expression, we relocated the congocidine antibiotic biosynthetic gene cluster (CGC) from its native terminal position to the central compartment in *Streptomyces ambofaciens*. This relocation enhanced CGC transcription compared to its original terminal location, both during exponential growth and after metabolic differentiation, resulting in 50% increase in congocidine production. At the 3D-level, transcription-induced domains formed at both the relocated and native CGC sites, creating sharp boundaries at a larger scale. Notably, the formation of such a boundary in the central compartment during the transition toward the stationary phase did not disrupt inter-arm contacts or affect neighboring gene expression. These results indicate that relocating a terminal cluster to the central chromosomal compartment provides a more favorable environment for transcription without altering chromosome compaction at the onset of metabolic differentiation, offering a promising strategy to enhance antibiotic production in the native host.

**Key points:**

- Central relocation of a gene cluster enhanced its transcription while preserving chromosome compaction.
- A transcription-induced domain formed at the new locus without altering neighboring gene expression.
- This strategy increased antibiotic yield by 50% in the native host.

## INTRODUCTION

*Streptomyces* bacteria are among the most abundant and ubiquitous soil-dwelling microbial genera (1). They are renowned for their prolific production of natural compounds, including antibiotics. *Streptomyces* produce around one third of all bioactive microbial metabolites discovered to date (2, 3). Despite the vast potential of specialized metabolite biosynthetic gene clusters (SMBGCs), with over 1 million sequences publicly available (4), only 3% of bacterial natural products have been identified to date (5). Consequently, the genomes of *Streptomyces* represent an untapped reservoir for the discovery of novel antibiotics.

*Streptomyces* are multicellular bacteria that undergo a complex life cycle, transitioning from unicellular spores to multicellular mycelia. (6–8). In these bacteria, antibiotic production is associated with metabolic differentiation and is typically induced by stress signals, such as nutrient depletion during the transition to the stationary phase, although production at low cell density has also been reported (for review, see (9)).Under starvation conditions, individual cells within the colony adopt distinct developmental fates, including antibiotic production, sporulation, programmed cell death, or dispersal to explore new environments (6–8). Antibiotic production is thought to contribute to colony survival by protecting nutrients released through cellular lysis or self-sacrifice, thereby benefiting the remaining cells.

Remarkably, *Streptomyces* possess a linear chromosome, an unusual feature among bacteria, characterized by terminal inverted repeats (TIRs) and specialized telomeres (10). Their chromosome is also relatively large (5–14 Mb) (11), GC-rich (∼72%) and genetically compartmentalized: the central region harbors highly conserved, essential genes and is transcriptionally active, while the arms are enriched in poorly conserved sequences, including SMBGCs (10). These clusters span from a few genes (*e.g.*, cyclodipeptide synthase or lanthipeptide clusters) to over a hundred kb in length (*e.g.*, stambomycin BGC). Within SMBGCs, genes undergo coordinated regulation. On average, each *Streptomyces* strain harbors around 30 SMBGCs (11), most of which remain silent under standard laboratory conditions. Even when SMBGCs are cloned for heterologous expression, this results in specialized metabolite production in only 11% to 32% of the cases (12, 13). Investigation into the causes of failure identified the lack of transcription as a major bottleneck, with no detectable transcripts in 60% of unsuccessful heterologous production attempts (12, 14). Understanding the molecular mechanisms governing the conditional expression of SMBGCs is therefore a key challenge to uncover the hidden metabolic potential of *Streptomyces*.

The three-dimensional architecture of the *Streptomyces* chromosome is intricately linked to its genetic compartmentalization and gene expression patterns during exponential growth (10). The central compartment, delimitated by distal ribosomal RNA (rrn) operons, exhibits distinct structural and transcriptional features compared to the terminal compartments (10). Notably, the central region consistently harbors the most highly expressed genes across species (15). This pattern holds irrespective of the degree of gene conservation or of the dose effect due to their proximity to the origin of replication (15). These observations suggest that the central compartment provides a transcriptionally permissive environment, contrasting with the lower transcriptional activity observed in the terminal regions. However, these findings are based on analyses of genes whose chromosomal positions have been shaped by evolutionary processes. The potential influence of compartmental location on gene expression remains to be explored.

To directly investigate the impact of chromosomal positioning on gene expression, we engineered the *Streptomyces ambofaciens* ATCC 23877 strain by relocating the 29 kb congocidine biosynthetic gene cluster (CGC) from its native terminal position to a central site within the chromosome. Congocidine (netropsin), a DNA minor groove-binding pyrrolamide, exhibits potent antimicrobial and antitumor activity. The CGC cluster encodes both biosynthetic enzymes, and the regulator Cgc1, which activates cluster expression during metabolic differentiation, when entering the stationary phase in response to an unidentified signal (16). Notably, congocidine further induces the expression of the gene *cgc*1 and of the resistance operon (including *cgc*20 and *cgc*21, encoding an ABC transporter), but does not affect other transcription units in the cluster, these remain activated only during the stationary phase (16).

Our results demonstrate that this targeted genomic engineering selectively enhances gene transcription and modulates chromosome architecture specifically at the relocated CGC locus, ultimately resulting in increased antibiotic production. These findings offer new insights into the regulatory mechanisms governing the conditional expression of SMBGCs, highlighting the critical role of the position within the chromosome in gene regulation.

## MATERIAL AND METHODS

### Bacterial strains, plasmids and experimental conditions

The bacterial strains and plasmids used in this study are listed in **Supplementary Table S1**. *Escherichia coli E*T12567/pUZ8002, carrying the plasmid of interest for intergeneric conjugation, was cultured in LB medium supplemented with kanamycin (25 µg/mL) and hygromycin (150 µg/mL on solid medium; 50 µg/mL in liquid medium). *Streptomyces s*trains were cultivated at 30°C on solid SFM medium (Soya Flour Mannitol: 20g/L organic soy flour, 20 g/L mannitol, 20 g/L agar) (17) for genetic manipulations and spore stock preparation.

The CGC cluster was inserted at the *att*B-PhiC31 site using a previously described conjugative and integrative cosmid carrying a 43.4 kb fragment encompassing the cluster (18) (**Supplementary Table S1**). This cosmid was derived from a genomic DNA library of *S. ambofaciens A*TCC23877 constructed in pBeloBAC11, into which a conjugative module and an *att-int P*hiC31 integrative system were subsequently inserted.

Plasmid conjugation from *E. coli i*nto *Streptomyces f*ollowed the method described by Kieser *et al. (*17). Exconjugants were selected on medium containing hygromycin (50 µg/mL) and nalidixic acid (50 µg/mL), and their genetic organization was confirmed by nanopore whole genome sequencing (Eurofins) **(Supplementary Data S1)**.

For analyses of congocidine production, RNA-seq, and Hi-C experiments, *Streptomyces* strains were grown in the congocidine production liquid medium MP5 (Medium of production n°5: 7 g/L yeast extract, 20.9 g/L MOPS, 5 g/L NaCl, 1 g/L NaNO_3_, 36 mL/L glycerol; pH 7.5) (19), as previously reported (20). Approximately 10^7^ spores were inoculated into 50 mL of liquid medium in a 500 mL Duran® Erlenmeyer baffled flask with a silicon stopper. Cultures were incubated at 30°C in a shaking orbital agitator (180 rpm, INFORS Unitron standard) and harvested after one day (Pseudo-opacimetry: OD_600nm_ ≈ 0.15, corresponding to ≈ 24 h of growth after inoculation; exponential phase) or two days (transition toward the stationary phase). In this medium, bacterial growth was monitored using pseudo-opacimetry (Biochrom WPA CO7500 colorimeter, Fisher Scientific). The term ‘pseudo-opacimetry’ reflects the limitations of this method due to the multicellular nature of these bacteria, which can affect accuracy. However, in MP5 medium, bacterial growth is sufficiently dispersed to allow reliable monitoring with this approach. To correct for minor differences in lag-phase duration between spore stocks, growth curves were normalized by setting the 24-h time point to a pseudo-OD of ∼0.15, based on a measured mean generation time of 3 h during exponential growth. The generation time during the exponential growth phase was determined from multiple pseudo-opacimetry measurements collected between OD 0.03 and OD 0.3 on Day 1. The measurements were fitted to the function OD600 = Ae^Bt^ , where *t* represents time and A and B are constants. The generation time was calculated as ln(2)/B. Culture aliquots were centrifuged and filtered using Whatman PVDF Mini-UniPrep syringeless filters (0.2 µm). Congocidine production was analyzed by HPLC as previously described (20).

### RNA preparation, sequencing and data processing

RNA was extracted from H1-CGC_native_-control and H1-CGC_reloc._ strains grown at Day 1 (OD_600nm_ ≈ 0.15) and Day 2 (24h later than the first sample, OD_600nm_ ≈ 2.8 ± 0.9 standard deviation, n = 14) accross seven independent experiments, then sequenced and analyzed as previously described (20). Reads were mapped to a reference genome containing a single TIR, with 7,170 gene features annotated according to *Streptomyces ambofaciens* ATCC23877 (GCF_001267885.1_ASM126788v1_genomic.gff, released 2025-04-26). This annotation was further refined to incorporate our manual curation of the CGC cluster. For differential gene expression analysis, the SARTools EdgeR-based R pipeline (21) was applied, using Day 1 growth of the ‘H1-CGC_native_-control’ strain (**Supplementary Table S1**) in MP5 medium as the reference condition, TMM default normalization and taking into account the batch effect. The RNA-seq analysis is presented in **Supplementary Table S3**. All data were analyzed using R software (22). The annotations in the ‘SMBGC’ column (**Supplementary Table S2**) were predicted using AntiSMASH (v8.0.4; (23)), with the exception of the CGC cluster, whose boundaries were manually curated.

### Marker frequency analysis (MFA) experiments

MFA was performed on four independent replicates of H1-CGC_native_-control and H1-CGC_reloc._ strains collected in parallel with RNA-seq samples. Pellets were extracted using phenol-chloroform, and DNA was sequenced using the Illumina NovaSeq platform with paired-end 150 bp reads. Reads were aligned to the same reference genome used for RNA-seq (containing only one TIR). Counts were generated for annotated genes, the CGC region (coordinates: 7,193,056–7,219,145 bp, excluding residual border sequences after *cgc* deletion at the native locus), an upstream region (‘CONGO_UP’, coordinates: 7,156,000–7,186,000 bp), and two 30 kb region around the origin of replication (‘ORI’, coordinates: 4,006,000-4,036,000 bp) and at the end (‘TER’, coordinates: 1-30,000 bp). Mapping was performed using Bowtie2, duplicates were removed with Picard, and counts were quantified using bedtools. Because the genome contains two TIRs, reads were mapped to a single TIR by combining the counts from both ends. Accordingly, the ORI/TER ratio for each sample was calculated by dividing the read count in the ORI region by half the read count in the TER region. The CONGO_UP/CGC ratio was calculated by dividing the counts in the CONGO_UP region by the counts in the CGC region for each sample. RNA-seq counts were normalized to MFA counts by converting to counts per million (CPM), computing the ratio of RNA-seq CPM to MFA CPM for each gene, and scaling by a factor of 200 to approximate original RNA-seq values. Normalized RNA counts were then analyzed using SARTools EdgeR as described above.

### Hi-C experiments

The Hi-C protocol, originally described by Cockram et al. (24), was adapted for *Streptomyces*. Bacteria were grown and collected under the same conditions as those used for the RNA-seq experiments. After one or two days of growth (strains listed in **Supplementary Table S1**), cultures were adjusted to a final OD_600_ of 0.15 in a volume of 89 mL. Proteins were cross-linked to DNA using 37–38% formaldehyde (Sigma Aldrich) at a final concentration of 3%, first for 30 minutes at room temperature (RT) with gentle agitation, followed by an additional 30 minutes at 4°C. Crosslinking was quenched with glycine (250 mM final concentration). Pellets were lysed in 600 µL of 1× TE buffer containing a protease inhibitor tablet and 8 µL (160 U) of Ready-Lyse lysozyme (LGC Biosearch, Cat# LU-R1804M), followed by incubation at RT for 45 min with gentle agitation (300 rpm, Eppendorf ThermoMixer™ C). Sonication was performed using a Bioruptor (Diagenode) with three cycles (30 seconds ON, 30 seconds OFF) on the low setting. SDS (10%) was added to each 300 µL sample (32.5 µL) and incubated for an additional 10 min at RT with gentle agitation (300 rpm, Eppendorf ThermoMixer™ C). For digestion, 600 µL of lysed cells were combined with a mix of 4,200 µL water, 600 µL NEB3 10× digestion buffer, and 600 µL Triton X-100 (10%). A 400 µL non-digested control was aliquoted, and 40 µL of SalI (20 U/ µL) was added to the sample for a 2h digestion at 37°C with agitation (180 rpm, INFORS Unitron standard). An additional 25 µL of SalI (20 U/ µL) was added, and incubation continued during 2 h under the same conditions. After digestion, 400 µL of the sample was reserved as a digestion control, and the remaining sample was centrifuged and resuspended in water.

Biotinylation was carried out for 1 h at 37°C with agitation (180 rpm, INFORS Unitron standard) using a mix of 13.5 µL dA/G/TTP (3.33 mM each), 50 µL 10× ligation buffer, 15 µL Biotin-14-dCTP (1 mM), 8 µL Large (Klenow) Fragment DNA Polymerase (40 U, NEB), and 13.5 µL water. Chromatin ligation was performed with a mix of 120 µL Thermo Scientific T4 10× ligation buffer, 12 µL BSA (10 mg/mL), 12 µL ATP (100 mM), 16 µL T4 DNA Ligase (30 U/µL), and 540 µL water. The mixture was incubated for 4 h at RT to allow ligation, followed by overnight incubation at 65°C with agitation (700 rpm, Eppendorf ThermoMixer™ C) to reverse cross-linking. Proteinase K, EDTA (6 mM final concentration) and SDS (0.5% final concentration) were then added, and samples were incubated for an additional 2 h at 65°C with agitation. DNA was purified with phenol:chloroform:isoamyl alcohol (25:24:1). Libraries were constructed as described in Lioy et al. (20), including DNA shearing (Covaris S220), end-repair/dA-tailing, adapter ligation, and library amplification. Sequencing was performed on an Illumina platform using the NextSeq 500/550 Mid Output Kit v2 for paired-end reads.

For Hi-C analysis, contact maps were generated using the OCHA pipeline in R (25) from paired-end sequencing data obtained after SalI restriction enzyme digestion. After mapping the reads of each strain, excluding TIRs and repetitive sequences from the cosmid, we removed PCR duplicates and invalid reads, including unmapped sequences. The remaining data were binned into sparse contact matrices and normalized using the balanced normalization method.

## RESULTS

### Impact of cluster relocalization on antibiotic production

To examine how genomic localization influences antibiotic production, we employed a previously described conjugative and integrative cosmid carrying a 43.4 kb fragment encompassing the cluster (18) (**Supplementary Table S1**). This construct was introduced into the *att*B-PhiC31 site of the S. ambofaciens H1-derived strain, CGCA018 (**Supplementary Table S1**), in which the native cluster had been deleted, yielding the recombinant strain ‘H1-CGC_reloc._’ (**Fig. 1.A**). In this strain, the CGC cluster retains the same orientation as in its native location relative to the origin of replication, with the majority of genes being transcribed in the same direction as the progression of the leading strand during replication (**Fig.1.B**). As a control, we used the parental strain ‘H1-CGC_native_-control’ (**Supplementary Table S1**), whichretains the congocidine gene cluster at its native locus and harbors at the *att*B-PhiC31 site a derivative of the pOSV806 integrative vector (26) devoid of reporter cassette. This control accounted for previously reported effects of PhiC31-mediated insertion on antibiotic production (27). Nanopore sequencing confirmed the correct single-copy insertion of the integrative vector into the target site for both the H1-CGC_native_-control and H1-CGC_reloc._ strains.

**Figure 1:**
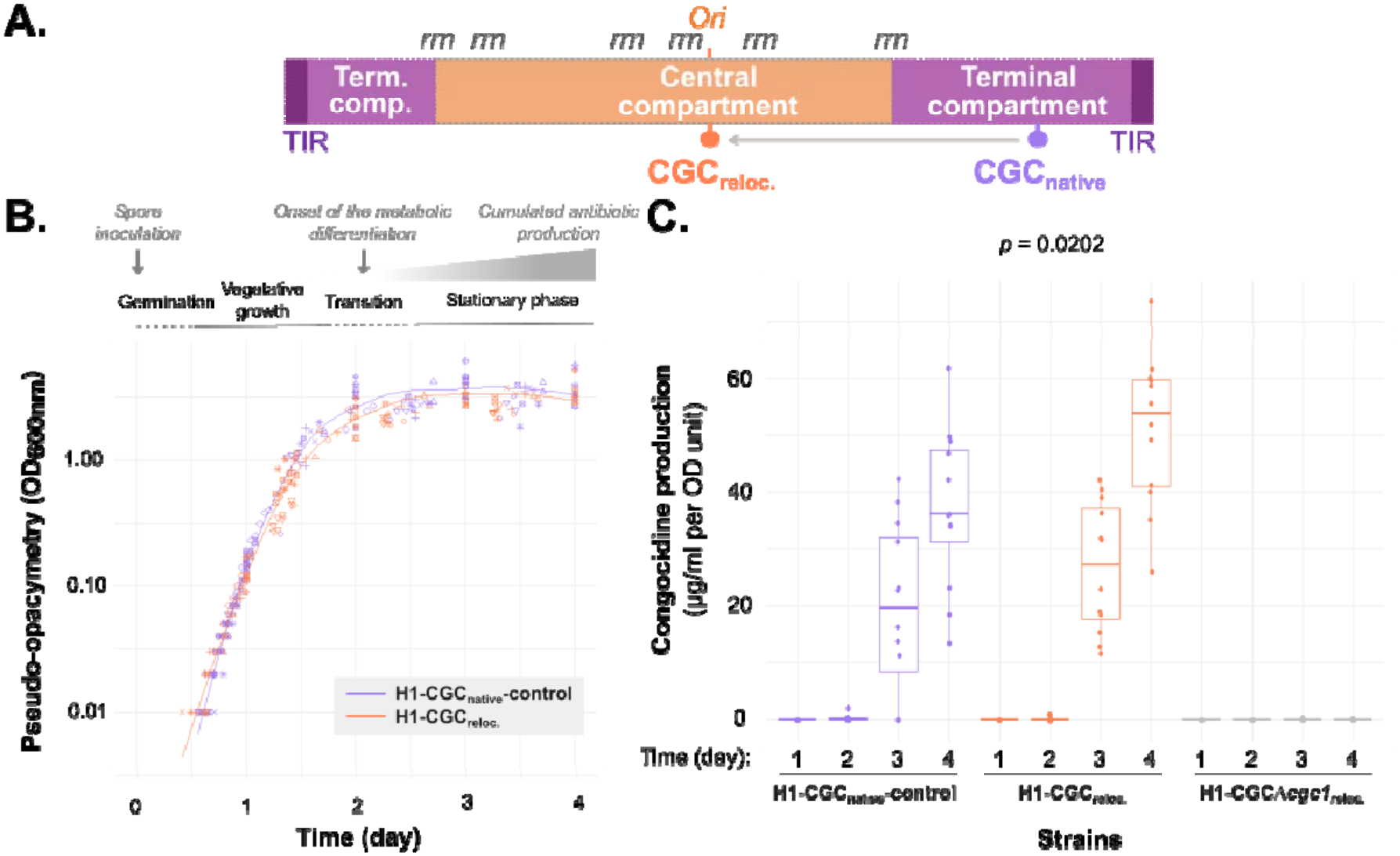
Growth and antibiotic production in strains with the CGC cluster at native or relocated positions. **A. Schematic representation of the genetic relocation of the CGC cluster (29 kb)from the terminal to the central chromosomal compartment.** Abbreviations: comp., compartment; *Ori*, origin of replication; *rrn*, ribosomal RNA operon; term., terminal; TIR, terminal inverted repeat. **B.Growth curves of H1-CGC**_**native**_**-control and H1-CGC**_**reloc**_. Around 10^7^ spores were inoculated into 50 mL of MP5 medium. Each point shape represents data from 14 independent experiments, each performed with two independent clones per strain. Different symbols represent independent biological experiments. A trend line was fitted using the LOESS (locally estimated scatterplot smoothing) method. **C.Congocidine production by strains harboring the CGC cluster at native or relocated positions**. Samples were collected during growth in MP5 medium. The strain H1-CGC_Δ_*cgc1*_reloc_. contains the CGC cluster at the relocated position but lacks the cgc1 gene encoding the cluster-situated regulator. This strain served as a negative control for congocidine production. Data were obtained from six independent experiments, each performed with two biological replicates. Statistical analysis was conducted using a Welch Two Sample *t-*test, with the p-value indicating the only significant difference between the native and relocated loci provided.

Both strains were cultivated in MP5 medium, which is optimized for antibiotic production (19). Growth monitoring revealed similar growth curves (**Fig. 1.B**). Both strains entered exponential growth one day after spore inoculation and exhibited similar generation times (**Fig. S1**). Consistently, the ORI/TER ratio decreased from 3.1 ± 0.1 SEM (n=8) on Day 1 to 2.0 ± 0.1 SEM (n=8) on Day 2, with no statistically significant difference observed between the two strains. Because the medium was not renewed during the experiment, we monitored the accumulation of antibiotic production over time. Detectable levels of congocidine production were observed from Day 3 onward. As commonly observed in *Streptomyces*, the developmental cycle is highly sensitive to environmental signals (9), which may account, at least in part, for the antibiotic production variability observed across the experiments presented here. Interestingly, by Day 4, during the late stationary phase, the median cumulative congocidine production was increased by 50% in the the H1-CGC_reloc._ Strain (median = 53.7, mean = 60.0, ± 3.8 µg.ml^-1^ per OD unit SEM, n=12) compared to H1-CGC_native_-control (median = 36.0, mean = 37.0, ± 4.0 µg.ml^-1^ per OD unit SEM, n=12) (**Fig. 1.C**).

The strain harboring a relocated CGC cluster devoid of the *cgc*1 gene encoding the cluster-situated regulator (‘H1-CGC_Δ_*cgc1*_reloc._’, **Supplementary Table S1**) fails to produce antibiotics (**Fig. 1.C**). This confirms that Cgc1 is essential for the biosynthesis induction (16). Importantly, this regulatory dependency persists even after genomic relocation, demonstrating that Cgc1 retains its native regulatory control over the cluster regardless of chromosomal position.

Taken together, these results demonstrate that relocating the cluster within its native host provides an additional strategy for increasing antibiotic production.

### Impact of cluster relocalization on its transcriptional activity

To further characterize the mechanisms underlying the increased antibiotic production after CGC relocation to the central compartment, we performed transcriptome and MFA analyses to assess its impact at both the RNA and DNA levels. Based on our prior observation that CGC cluster transcription is already induced by Day 2 (16, 20), we conducted experiments at two growth stages: the exponential phase (Day 1) and the transition to the stationary phase (Day 2), corresponding to the onset of metabolic differentiation. The latter condition was selected to capture the early stages of CGC cluster induction and to ensure higher-quality RNA preparations, as RNA quality deteriorates over time of cultivation in our experience.

As shown in **Figure 2**, during the exponential phase, we observed a mild increase in the expression of *cgc* genes following CGC relocation to the central compartment. This increase was observed across all *cgc* genes, with an average fold change of 2.6 (± 0.1 SEM, n=22) when considering a p-value < 0.05, and for 12 of the 22 genes, with an average fold change of 2.9 (± 0.1 SEM, n=12) when considering an adjusted p-value < 0.05. To determine whether this increase could be attributed to differences in CGC copy number, we used MFA to quantify the number of CGC cluster copies relative to a reference sequence of similar size (30 kb) located upstream of the cluster at the native site (positions: 7,156,000 – 7,186,000 bp). This analysis was performed for four of the seven replicates. We found that the CGC copy number was approximately 2.1-fold higher (± 0.06 SEM, n=4) in the H1-CGC_reloc._ strain than in the H1-CGC_native_-control strain, which could completely account for the increased *cgc* gene expression observed during this phase. Consistent with this interpretation, normalization of the expression data according to CGC copy number abolished the observed statistical differences (**Table S2**). Together, these results suggest that relocation of the CGC cluster increases its basal transcriptional level during the exponential phase through a gene dosage effect.

**Figure 2:**
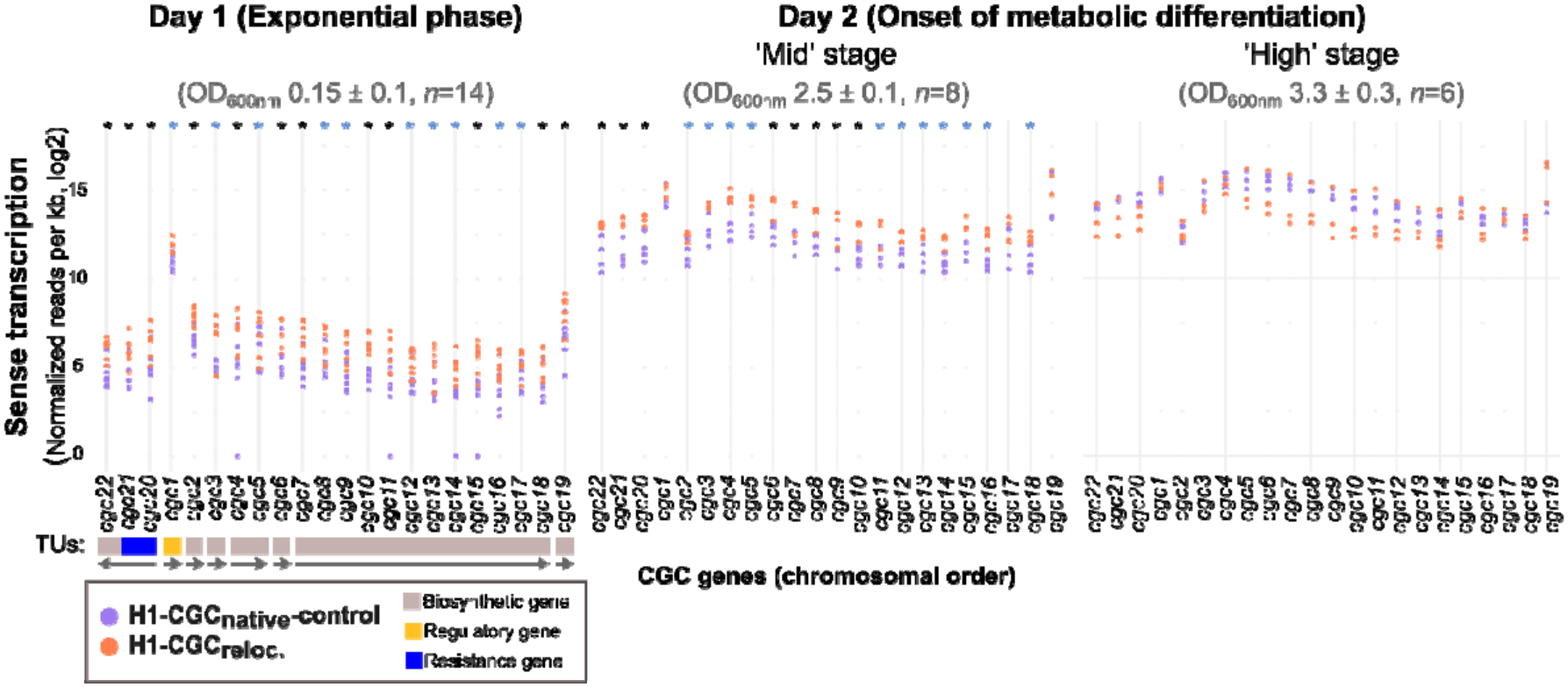
Dynamics of the congocidine gene cluster transcriptome across growth phases and chromosomal positions. H1-CGC_reloc._ and H1-CGC_native_-strains were grown in liquid MP5 medium for two days in seven independent experiments. On Day 2, biological replicates were classified as ‘Mid’ or ‘High’ metabolic differentiation stages based on the level of CGC expression in the H1-CGC_native_ strain. For four of these replicates, MFA experiments were performed to quantify the CGC copy number relative to a reference region located upstream of the native CGC cluster. The CGC-to-reference ratio was 2.1 ± 0.1 SEM (*n* = 4) and 1.4 ± 0.1 SEM (*n* = 4) on Days 1 and 2, respectively, with no statistically significant difference between the ‘Mid’ and ‘High’ stages on Day 2. Each data point represents normalized read counts obtained using EdgeR and adjusted for gene length, from independent RNA-seq experiments (seven in the exponential phase, four at the ‘Mid’ stage, and three at the ‘High’ stage). Data are shown in orange for the H1-CGCreloc. strain and in violet for the H1-CGCnative-control strain. Genes showing statistically significant induction (adjusted p-value ≤ 0.05) are marked with a black asterisk. Genes with an unadjusted p-value ≤ 0.05 but an adjusted p-value > 0.05 are marked with a blue asterisk. The eight previously characterized transcription units (TUs) (16) are indicated by rectangles below the gene names, with colors reflecting their encoded functions and arrows indicating the direction of transcription. Genes are presented in their chromosomal order, with the left side corresponding to the region closer to the origin and the right side to the region closer to the chromosome end.

We further analyzed the impact of CGC relocation at the onset of the metabolic differentiation (Day 2, **Fig.2**). Since Day 2 represents a transitional phase, we were able to collect samples at slightly distinct stages of metabolic differentiation. We conducted seven independent experiments, which separated into two patterns based on the level of CGC induction at the native locus, correlating with the pseudo-opacimetry measured at Day 2: When the CGC is located at its native position, we observed mid-induction of the CGC cluster at the lowest OD (around 2.5, hereafter referred to as the ‘Mid’ condition), compared to higher induction at higher OD (above 3, hereafter referred to as the ‘High’ condition). This observation aligns with previous findings that increased cell density induction often correlates with higher antibiotic production, a process that may involve quorum sensing (9). Therefore, we analyzed these two induction levels separately. We observed that the copy number of the relocated CGC was similar under both the ‘Mid’ and ‘High’ conditions, being 1.4-fold higher (± 0.1 SEM, *n =* 4) than that of the CGC at the native position.

In the ‘Mid’ condition, all *cgc g*enes, except *cgc1, cgc17 a*nd *cgc19*, were overexpressed in the relocated strain when considering a *p-v*alue < 0.05 (fold change of 3.1 ± 0.1 SEM, *n=*19), with eight of them showing statistically significant induction with an adjusted *p-*value < 0.05 (fold change of 3.5 ± 0.1 SEM, *n=*8) (Fig.2). In this condition, the dose effect alone, corresponding to a 1.4-fold increase, does not fully account for the observed increase in expression following CGC relocation.

In the ‘High’ condition, we no longer observed statistically significant differences in CGC gene expression between the H1-CGC_reloc._ and H1-CGC_native_-control strains (**Fig.2**), despite the copy number being 1.4-fold higher at the relocated position. This suggests that CGC expression may have reached a plateau at this stage.

Overall, these results indicate that relocation of the CGC cluster to the *att*B-PhiC31 site enables the strain to reach the high-expression state earlier. Moreover, the increase in CGC gene expression under the exponential and ‘Mid’ conditions was approximately 2–3-fold, exceeding the 50% increase observed in antibiotic production (**Fig.1.C**). This discrepancy suggests that post-transcriptional processes may represent rate-limiting steps in congocidine biosynthesis, limiting the increase in antibiotic production despite higher transcript levels.

### Impact of cluster relocalization on chromosome folding

We have previously demonstrated that during metabolic differentiation, the chromosome adopts a more compact conformation, characterized by the emergence of a secondary diagonal on contact maps (20). Notably, the CGC cluster forms a distinct boundary within the terminal compartment (20). This observation led us to hypothesize that the evolutionary enrichment of SMBGCs at chromosome ends may serve to minimize the disruptive impact of their strong induction on the formation of this secondary diagonal, a process driven by SMC-condensin activity, as previously demonstrated during sporulation in *Streptomyces venezuelae* (28).

To test this hypothesis, we examined the effects of genetic relocalization of the congocidine cluster on chromosome conformation. As shown in **Figure 3**, relocalization had minimal impact during the exponential phase, with no detectable large-scale structural changes. However, following metabolic differentiation, a sharp boundary emerged at both the native and relocated sites, correlating with high CGC cluster expression. By focusing on this CGC-associated boundary, we can observe the formation of a transcription-induced domain (TID) (**Fig. 3**), a phenomenon previously described in *E. coli* using artificial systems (29) and in natural systems such as the Salmonella pathogenicity island (30). Importantly, the secondary diagonal remained visible in both strains (**Fig. 3**).

**Figure 3:**
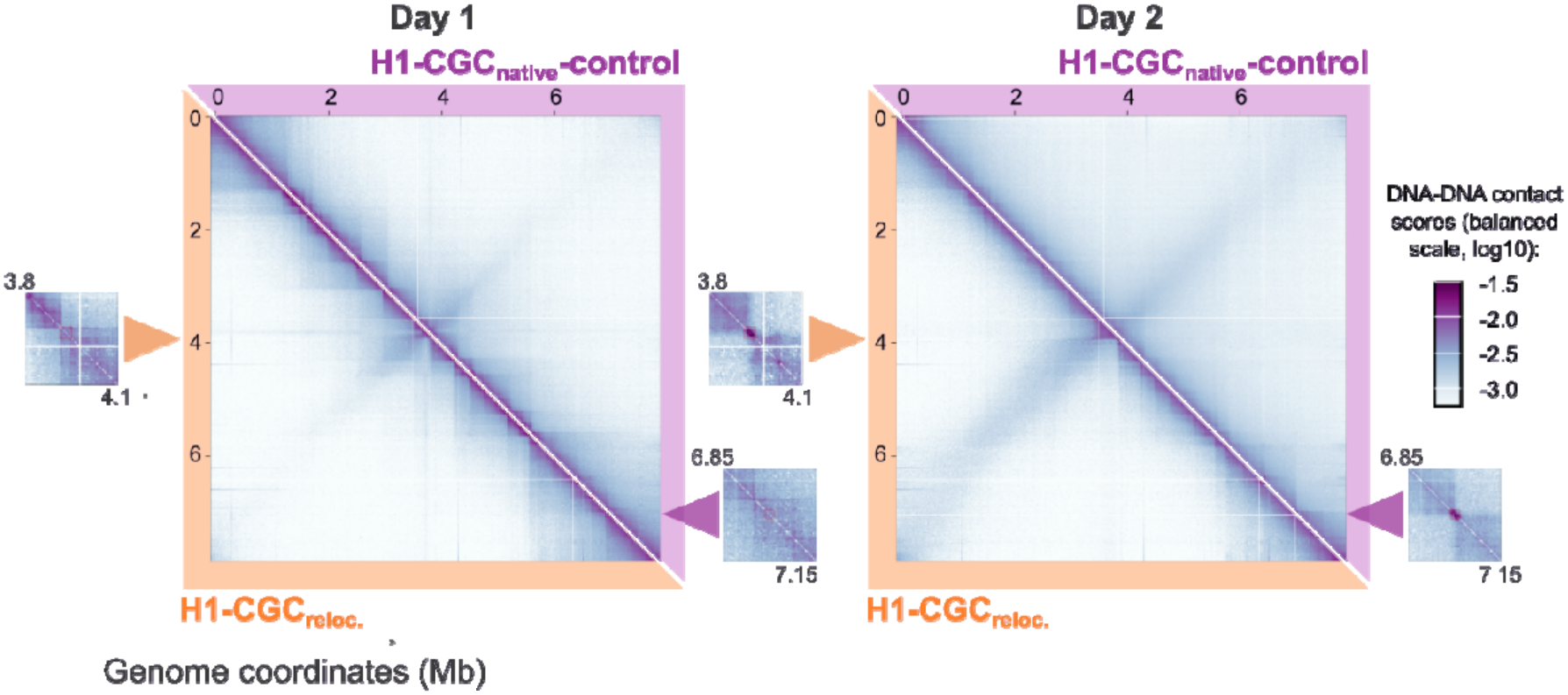
Dynamics of chromosome spatial conformation in strains harboring the CGC cluster at native or relocated positions. Hi-C experiments were performed on H1-CGC_native_-control (violet) and H1-CGC_reloc._ (orange) strains grown in MP5 medium during exponential phase (Day 1) or during transition toward the stationary phase (Day 2). The normalized contact map at 10 kb resolution was generated from asynchronous populations, with the x and y axes representing genomic coordinates after excluding TIRs. The color scale indicates contact frequency between genomic loci, ranging from white (rare contacts) to dark purple (frequent contacts). Inset panels provide a zoomed-in view of a 300 kb region at 5 kb resolution, highlighting the CGC cluster (red square), with genomic positions indicated in megabases (Mb). The data represent the merged results of two to four independent experiments, while individual replicates and their quantitative characteristics are detailed in **Supplementary Figure S2**. White lines across all maps correspond to regions with low SalI site density, as shown in **Supplementary Figure S3**. Additional zoomed-in views of a 300 kb region, including the *attB*-PhiC31 site (20 kb red box), are presented for the parental strain without insertion and the H1-CGC_native_-control strain in **Supplementary Figure S4**. The observed/expected maps of the Hi-C matrices are provided in **Supplementary Figure S5**.

These findings suggest that cluster relocalization does not disrupt the dynamic properties of the chromosome, including the interarm interactions characteristic of genome compaction during the early stationary phase, and the formation of a TID at the CGC cluster, which underlies the sharp boundary.

### Impact of cluster relocalization on gene expression at the integration site and genome-wide host responses

Given the significant transcriptional effects resulting from relocating the CGC cluster to the *att*B-PhiC31 site, we investigated whether this genomic modification altered the expression of genes flanking the insertion site. Surprisingly, we detected very limited differential gene expression in the *att*B-PhiC31 region between strains harboring the CGC insertion and those containing only the control vector, at both Day 1 and Day2 (**Fig. 4**). Remarkably, the formation of a sharp 3D-boundary at the insertion site does not disrupt transcription of genes located in adjacent chromosomal interacting domains (CIDs), a finding that may challenge initial expectations.

**Figure 4:**
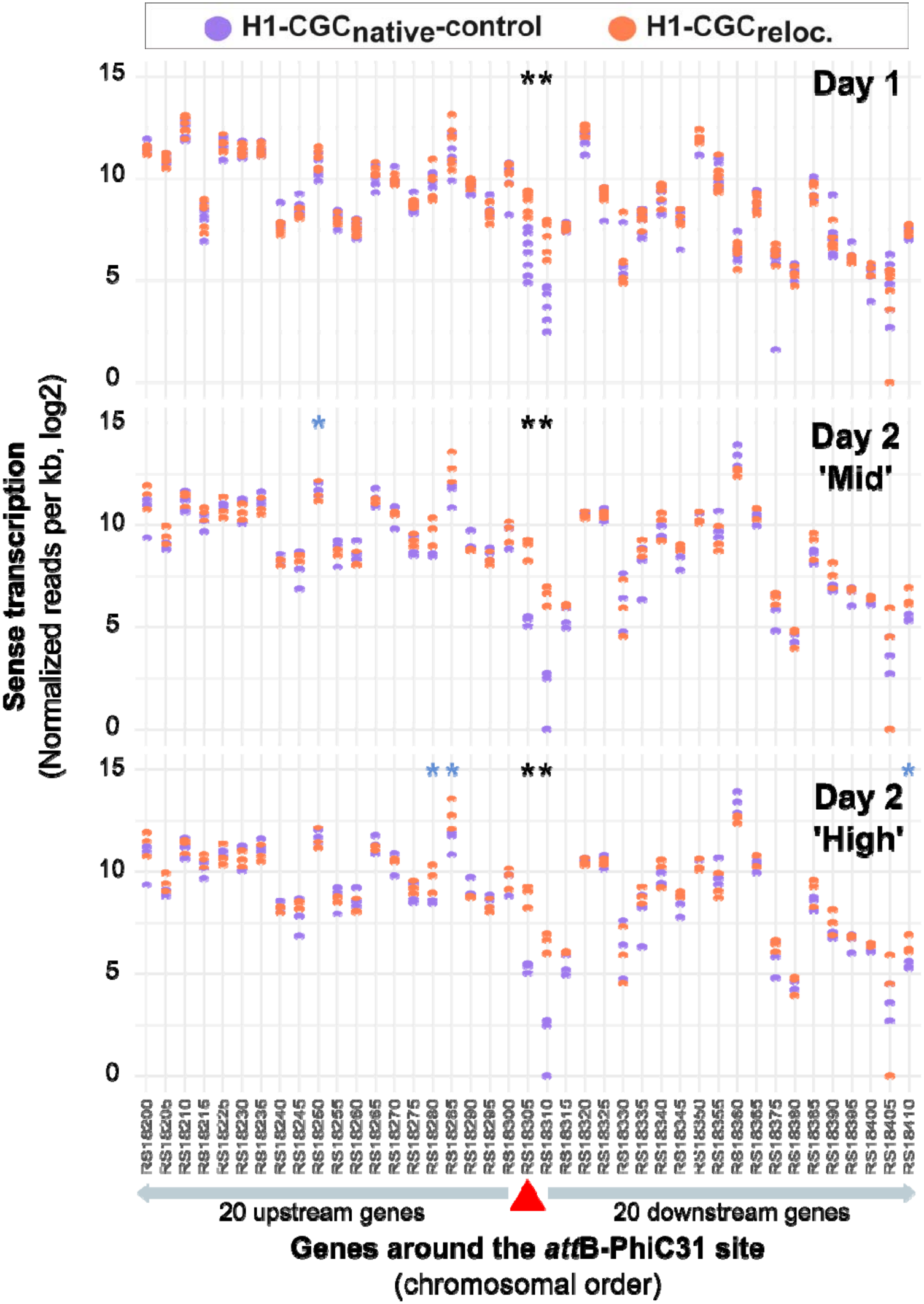
Sense transcriptome profiles around *att*B**-PhiC31 site in strains harboring the CGC cluster at native or relocated positions**. RNA-seq experiments were performed using MP5 growth medium, with samples collected during both the exponential phase (Day 1) and during the transition toward the stationary phase at ‘Mid’ or ‘High’ stages (Day 2). Each data point represents normalized read counts, processed using EdgeR and adjusted for gene length, for 20 genes located upstream and downstream of the *attB-*PhiC31 target site (*SAM23877_RS18305*, red triangle), based on seven independent RNA-seq experiments (as described in the Fig.2 legend). Gene names correspond to the *SAM23877_RS a*nnotation. *SAM23877_RS18370 w*as excluded from the analysis because no reads were detected for this gene.. Data for the H1-CGC_reloc._ strain are shown in orange, whereas data for the H1-CGC_native_-control strain are shown in violet. Genes showing statistically significant induction (adjusted *p-*value ≤ 0.05) are marked with a black asterisk. Genes with an unadjusted *p-*value ≤ 0.05 but an adjusted *p-*value > 0.05 are marked with a blue asterisk. The expression levels of both the gene into which the CGC cluster was inserted and the downstream gene were significantly increased in the relocated strain compared with the native control strain across all conditions tested. The increase ranged from 5- to 12-fold for the former and from 9- to 24-fold for the latter, depending on the experimental conditions (**Supplementary Table S3**)..

Finally, genome-wide analysis revealed limited differential expression of genes outside the main cluster when comparing the two strains, affecting only 0.5–1% of all genes when comparing the two strains (**Supplementary Fig. S6, Supplementary Table S3**). Among the genes consistently differentially expressed (adjusted *p-*value < 0.05), we identified the PhiC31 target site and its downstream gene (**Fig. 4**), three genes located within or in the vicinity of the Samy prophage, three genes flanking the CGC cluster within the vector used for strain engineering, and two genes 150 kb downstream of the CGC cluster (*SAM23877_RS32470 a*nd *SAM23877_RS32480)* (**Supplementary Table S3**). Some genes belonging to a RiPP-like cluster (Ribosomally synthesized and Post-translationally modified Peptide; position: 7,314,194-7,324,409bp) were also found to be downregulated in the relocalized strain compared with the native strain These changes in gene expression are limited and may reflect adaptive responses, potentially triggered by the physiological stress associated with CGC overexpression during the early developmental stage, and/or the genetic modifications. Indeed, the genome engineering process for this strain involved a series of challenging steps: first, the removal of the CGC cluster from its native locus, followed by the excision of the selection marker (**Supplementary Table S1**), and finally, the introduction of an integrative cosmid carrying the CGC cluster.

Collectively, these findings indicate that relocating the CGC cluster has minimal impact on the expression of genes surrounding the integration site. Thus, at the local level, the observed increase in CGC gene transcription appears to be specific to the cluster and does not extend to neighboring genes at the PhiC31 integration site. The genome engineering of the CGC cluster, which is the most highly expressed SMBGC during the early stationary phase (**Supplementary Fig. S6, Supplementary Table S3**), may have broader physiological implications beyond its direct transcriptional regulation, which remains limited to at most 1% of all genes.

## DISCUSSION

The limited expression of SMBGCs remains a critical bottleneck in natural product research, primarily due to transcriptional constraints (14, 31). To overcome this challenge, multiple strategies have been developed, including heterologous expression, promoter engineering, regulatory manipulation through the overexpression or deletion of cognate regulators (31), and the application of transcription factor decoys (32). However, the heterologous production of natural products is often strongly influenced by host-specific regulatory and biosynthetic factors, making chassis selection largely empirical. Although phylogenetically related hosts are generally considered more suitable for expressing foreign BGCs, this assumption does not universally hold true (33). In this study, we extend current strategies for enhancing SMBGC expression by investigating how chromosomal positioning affects SMBGC activity within the native host.

Specifically, we show that relocating the CGC cluster to the *att*B-PhiC31 site significantly enhances its transcriptional activity in the native host. Given that heterologous integration of SMBGCs frequently occurs at the *att*B-PhiC31 site, these findings suggest that, in some cases, the apparent benefits of heterologous expression may arise from the genomic integration site itself rather than from the choice of chassis. This locus is widely used for genetic insertions, and previous studies have reported that integration at this position can influence antibiotic production (27, 34). To minimize potential positional artifacts, we included an internal control (pOSV806_Δ_; **Supplementary Table S1**). A broader, systematic analysis of alternative chromosomal loci could further identify genomic positions that are optimal for both native and heterologous SMBGC expression.

While position effects on gene expression have been documented in heterologous systems (35–37), to our knowledge this is the first study to apply this concept to autologous SMBGC expression. Our findings suggest that SMBGCs are not necessarily positioned for maximal transcriptional activity within their native chromosomal context. Instead, their localization may act as a regulatory constraint that modulates or even limits cluster expression. This hypothesis warrants broader investigation across different SMBGCs and integration sites.

Several, non-mutually exclusive mechanisms could explain the increased transcription observed following relocation of the CGC cluster to the *att*B-PhiC31 site, both during the exponential phase and at the early stage of the transition to the stationary phase (Day 2, ‘Mid’ condition). Increased expression could result from (i) an increased copy number of the cluster, leading to a gene dosage effect; (ii) a more transcriptionally permissive molecular environment in the central chromosomal compartment compared with the native location of the cluster; and/or (iii) a greater responsiveness of the CGC cluster to metabolic differentiation occurring during the transition to stationary phase, as its higher basal transcription during exponential phase may prime or accelerate the subsequent induction of the cluster. Below, we discuss these three hypotheses in turn. During the exponential phase, we observed that relocation of the CGC cluster to the central chromosomal position was associated with a higher basal transcription, which can be largely explained by the approximately twofold higher copy number of the cluster at the central position compared with its native location (**Table S2**). On Day 2, however, the CGC cluster copy number was only 1.4-fold higher than at its native position and therefore appears insufficient to fully account for the approximately threefold increase in CGC expression. Although gene dosage may account for a substantial proportion of the transcriptional effects associated with gene position in bacteria (38), other mechanisms may also contribute. For instance, relocating the fis gene from an oriC-proximal to a terminus-proximal position in E. coli caused a fitness defect despite unchanged fis transcript and FIS protein levels, suggesting that positional effects independent of copy number may arise from the limited diffusibility of RNA and proteins, thereby altering the activity of the FIS regulon (39). Thus, the additional increase in CGC expression observed following relocation may reflect distinct transcriptional properties of the central and terminal chromosomal compartments and/or more efficient activation of the CGC cluster during the transition to stationary phase. Supporting the hypothesis of distinct molecular properties of the central and terminal compartments, chromosomal organizers and nucleoid-associated proteins, such as HupA and SMC condensin, are enriched in the central region, while HupS predominates in the terminal compartments (28, 40). These compartments likely exhibit divergent chromatin composition and transcriptional activity, analogous to the A/B compartments described in mammalian genomes (41, 42) and certain archaea (43). Notably, in *Escherichia coli*, regions near rrn operons display higher transcriptional propensity (44). Under nutrient-rich conditions, RNA polymerase forms dense clusters engaged in ribosomal RNA synthesis (45–47), suggesting that central regions may be enriched in transcription and translation machinery, thereby favoring active gene expression. Consistent with this model, we previously reported in *Streptomyces* species that genes located in the central compartment are more highly expressed than those in the terminal regions (15). Our current study corroborates this pattern using the same gene set, the CGC cluster, further reinforcing the concept of functional chromosomal compartmentalization in bacteria. Finally, regarding the last hypothesis, basal transcription levels can often prime SMBGC activation, such that even modest increases in expression may be sufficient to trigger clusters controlled by self-amplifying regulatory loops. This is illustrated by the positive feedback mechanism of congocidine antibiotic on the CGC cluster (16). This hypothesis could explain why relocation primarily results in the earlier attainment of high expression levels, which appear to plateau under the conditions tested (**Fig.2**).

Interestingly, we identified promoters within the CGC cluster that exhibited distinct responses to relocation in the central genomic region at the onset of metabolic differentiation, under the ‘Mid’ condition. Specifically, P_*cgc*1_, the promoter driving expression of the cluster-situated regulator Cgc1, and P_*cgc*19_ were not significantly upregulated in the relocated strain under this condition. Notably, these two promoters also exhibit the highest transcriptional activity within the cluster. These differences may reflect variations in RNA polymerase affinity or local promoter architecture. This raises an intriguing possibility: that SMBGCs may possess a regulatory architecture combining robust, localization-insensitive expression of key regulators with localization-sensitive expression of biosynthetic genes. Further analyses across diverse clusters are needed to test this hypothesis.

The mechanisms underlying position-dependent regulation likely involve multiple spatial and molecular factors, including RNA polymerase availability, reduced efficacy of Lsr2-mediated silencing, local DNA supercoiling, promoter strength, and connectivity with regulatory networks.

Despite relocation, the CGC cluster retained its native regulatory behavior, including stationary-phase induction under Cgc1 control. Although congocidine activates the transcription of *cgc1* and efflux genes, the exact signal for full cluster induction by this orphan response regulator remains unknown (16). Notably, relocation enhanced both transcription and antibiotic production, although the latter to a lesser degree. This discrepancy suggests that transcription is no longer the primary limiting factor; rather, post-transcriptional processes, such as translation efficiency, protein folding, metabolic flux or availability of phosphopantetheine transferases (48) or MbtH-like partners (49), may now limit production. Thus, chromosomal repositioning effectively relieved the transcriptional bottleneck, though its generalizability to silent clusters remains to be determined.

By analyzing nearly one thousand bacterial genomes, Hu et al. (50) showed that approximately two-thirds of bacterial gene families exhibit positional biases in bacteria with circular chromosomes, with genes preferentially localized near either the origin or the terminus of replication, particularly in fast-growing species. In the linear genome of *Streptomyces*, this phenomenon is particularly pronounced and is referred to as ‘genetic compartmentalization’. This positional bias also extends to SMBGCs, which display family-specific enrichment toward the terminal regions of the chromosome (51). In addition to hypotheses involving gene dosage or other transcriptional effects, as previously proposed in other bacteria (38), one might speculate that the terminal localization of SMBGCs has evolved to help preserve overall genome compaction during the transition to stationary phase. Our results show that the formation of a sharp boundary associated with SMBGC induction near the origin of replication does not disrupt the secondary diagonal observed in DNA contact maps, which reflects large-scale chromosomal interactions. Moreover, our data suggest that transcriptional activity can shape 3D chromosomal conformation but not vice versa, at least at the *att*-PhiC31 site. In eukaryotes, disrupting topological domains (TADs), which are analogous to bacterial CIDs, can typically alter gene expression (52). However, in our study, focused on the impact of CGC relocation to the *att*B-PhiC31 site, we observed no such effect. Specifically, the formation of a sharp boundary at this locus did not markedly influence the transcription of adjacent genes. These results highlight the unidirectional influence of transcription on local chromosomal architecture in *Streptomyces*, emphasizing the complexity of spatial genome organization and its regulatory implications. Taken together, our results rather support the hypothesis that the terminal localization of SMBGCs may have been selected, at least in part, for transcriptional reasons, helping to prevent excessive basal transcription during the exponential phase and, more broadly, to regulate the timing of their induction. Finally, beyond demonstrating the formation of a barrier upon CGC cluster activation, the data presented in Figures 3 and 4 demonstrate that cluster relocation does not perturb overall chromosome organization or induce widespread transcriptional changes.

Overall, this study introduces position within the chromosome as a powerful approach to modulate SMBGC expression in *Streptomyces*. By integrating chromosome positioning into the regulatory framework of specialized metabolism, our findings provide a new perspective for optimizing both native and heterologous production of natural products.

## DATA AVAILABILITY

The RNA-seq and HiC-seq data generated during this study have been deposited in the European Nucleotide Archive (ENA), under the project accession number PRJEB108791 (ERP189626).

## SUPPLEMENTARY DATA

Supplementary Data are available at NAR online.

## AUTHOR CONTRIBUTIONS

SBM: Conceptualization, Formal analysis, Hi-C and RNA-seq experiments, Writing - original draft. SL, JLP: Conceptualization, Writing - review & editing. HJ: Hi-C experiments, Writing - review & editing. ND & TG: Phenotypic characterization of the strains (growth, HPLC analyses), Writing - review & editing. HL: Obtention of the mutant strains, Writing - review & editing.

## ACKNOWLEDGEMENTS

We thank Corinne Saulnier for her technical help. We acknowledge the sequencing and bioinformatics expertise of the I2BC High-throughput sequencing facility, supported by *France Génomique (*funded by the French National Program *Investissement d’Avenir A*NR-10-INBS-09). We also thank Pascaline Tirand and Fanny Culot for their daily help.

