## Supplementary information for "Relocating the congocidine biosynthetic cluster to the central chromosomal compartment enhances gene expression and antibiotic production in its native host, *Streptomyces ambofaciens*"

### Supplementary Figures and Tables

**Supplementary Figure S1**: Comparison of the growth of strains H1-CGC$_{native}$-control and H1-CGC$_{reloc.}$ in MP5 liquid medium

**Supplementary Figure S2**: Individual Hi-C contact maps underlying the merged maps presented in **Figure 3**

**Supplementary Figure S3**: Distribution of SalI Sites along the chromosomes of H1-CGC$_{native}$-control and H1-CGC$_{reloc.}$ strains

**Supplementary Figure S4**: Zoomed-in views of the *attB*-PhiC31 site in the parental strain without insertion and the H1-CGC$_{native}$-control strain

**Supplementary Figure S5:** Observed/Expected (OE) maps of the Hi-C matrices presented in Figure 3

**Supplementary Figure S6:** MA plot of differential gene expression in H1-CGC$_{reloc.}$ and H1-CGC$_{native}$ control strains over growth

**Supplementary Table S1**: Bacterial strains and plasmids used in this study

**Supplementary Table S2:** Fold change in *cgc* gene expression during exponential phase (Day 1) in the H1-CGC$_{reloc.}$ and H1-CGC$_{native}$-control strains, before and after normalization to MFA counts

**To download on line:**

**Supplementary Table S3**: Results from the RNAseq approach conducted in this study. The legend is detailed in the "Readme" sheet.

**Supplementary Data 1**: Nanopore sequencing results (Eurofins) for *S. ambofaciens* ATCC 23877 strains H1-CGC$_{native}$-control and H1-CGC$_{reloc.}$ (W23 and X24 clones)

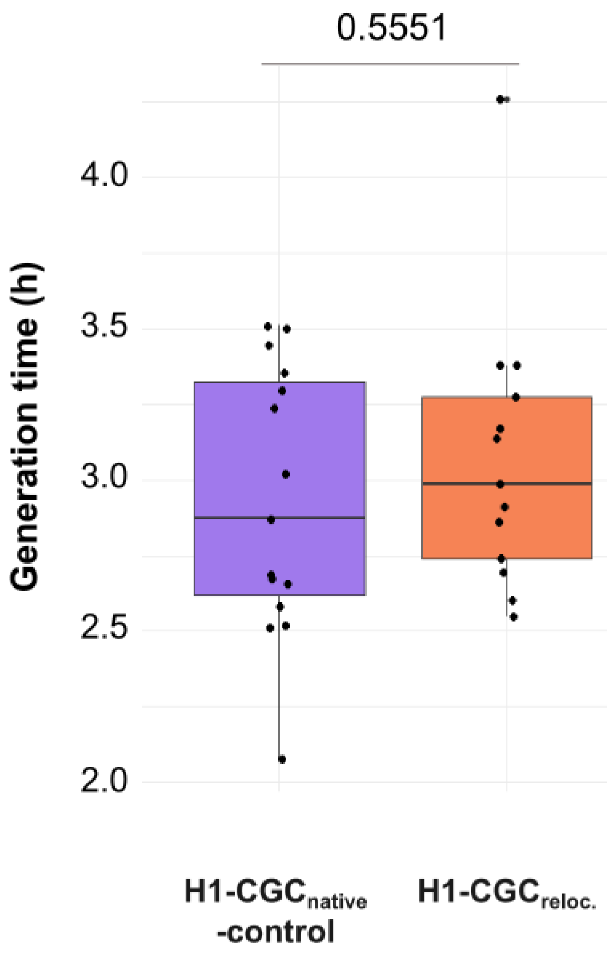


**Figure S1: Comparison of the generation times of strains H1-CGC$_{native}$-control and H1-CGC$_{reloc.}$ during the exponential growth phase (Day 1) in MP5 liquid medium**

Results are derived from 8 independent experiments, each performed with two independent clones per strain. Statistical significance was assessed using an exact Wilcoxon rank-sum test, and the corresponding *p*-value is reported.

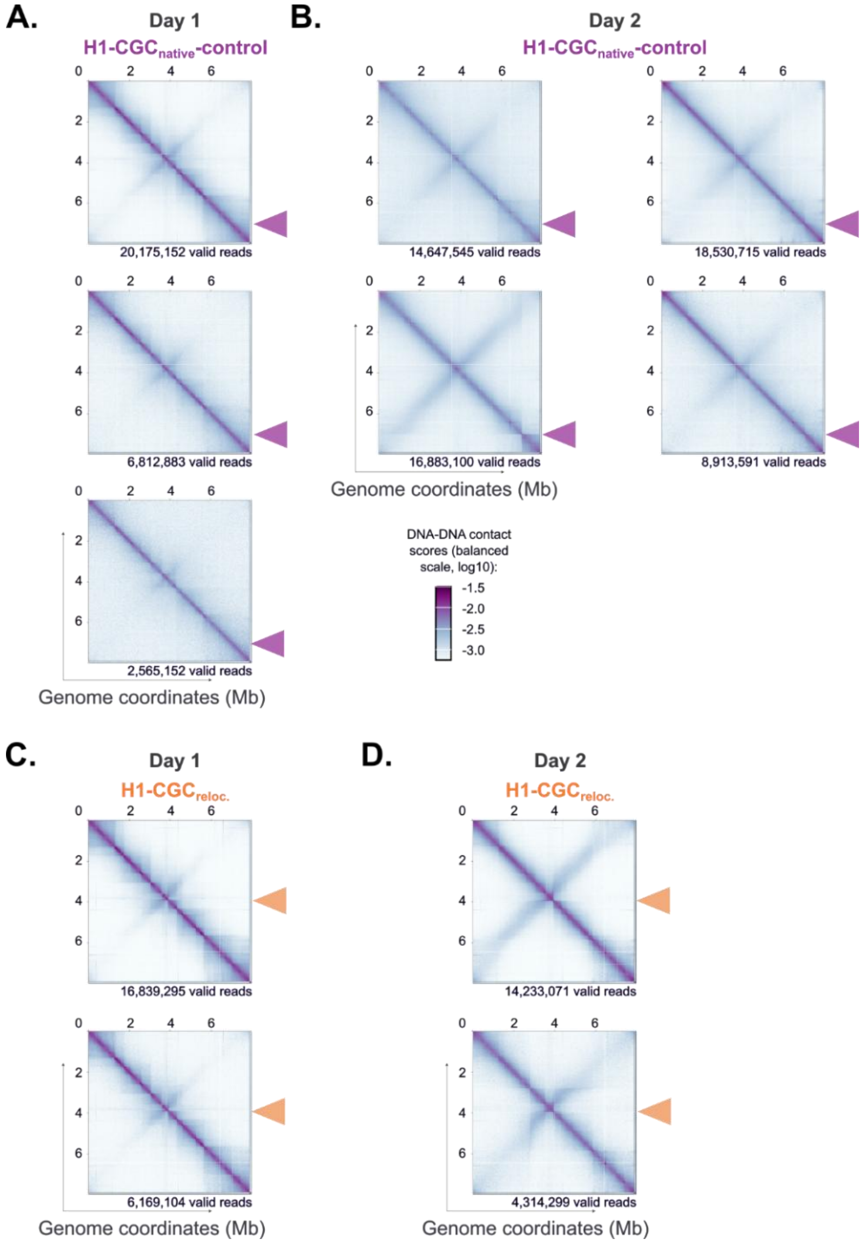


**Figure S2: Individual Hi-C contact maps underlying the merged maps presented in Figure 3**

Hi-C experiments were performed on H1-CGC$_{native}$-control (violet, A and B panels) and H1-CGC$_{reloc.}$ (orange, C and D panels) strains grown in MP5 medium. Data were collected during two growth phases: the exponential phase (Day 1, A and C panels) and the transition toward stationary phase (Day 2, B and D panels). The normalized contact maps, displayed at a 10 kb resolution, were generated from asynchronous cell populations. The x and *y* axes represent genomic coordinates, excluding terminal inverted repeats (TIRs). The color scale reflects the frequency of contacts between genomic loci, ranging from white (rare contacts) to dark purple (frequent contacts). This scale is consistent across all Hi-C maps and is depicted in Panel B. The number of valid reads for each matrix is indicated below it. The interaction results for plasmid pSAM1 are displayed in the contact maps to the right of the solid black line. The triangles indicate the position of the CGC cluster in each strain. Please note that a boundary at the desferrioxamine cluster (located at 2.8 Mb, excluding the first TIR) can also appear in some Hi-C matrices of *S. ambofaciens* obtained at Day 2, as shown in one of the replicates in panel B.

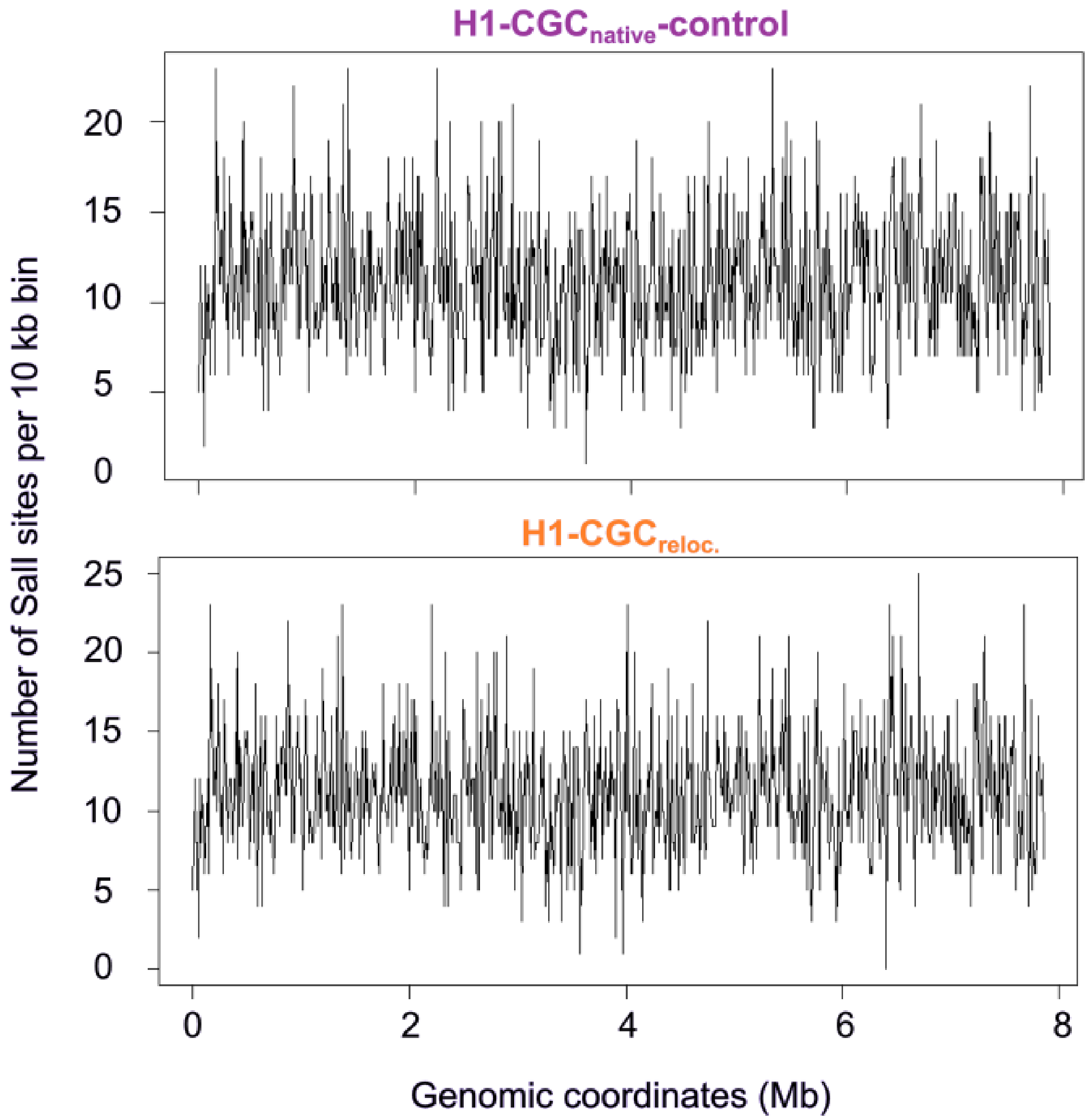


**Figure S3: Distribution of SalI Sites along the chromosomes of H1-CGC$_{native}$-control and H1-CGC$_{reloc.}$ strains**

The number of SalI restriction sites was counted in 10 kb windows (10 kb step size) along the chromosomes of H1-CGC$_{native}$-control and H1-CGC$_{reloc.}$ strains, excluding TIRs. Regions with the lowest site density appear as white lines on some Hi-C maps, reflecting poor coverage due to limitations in the Hi-C data generation process.

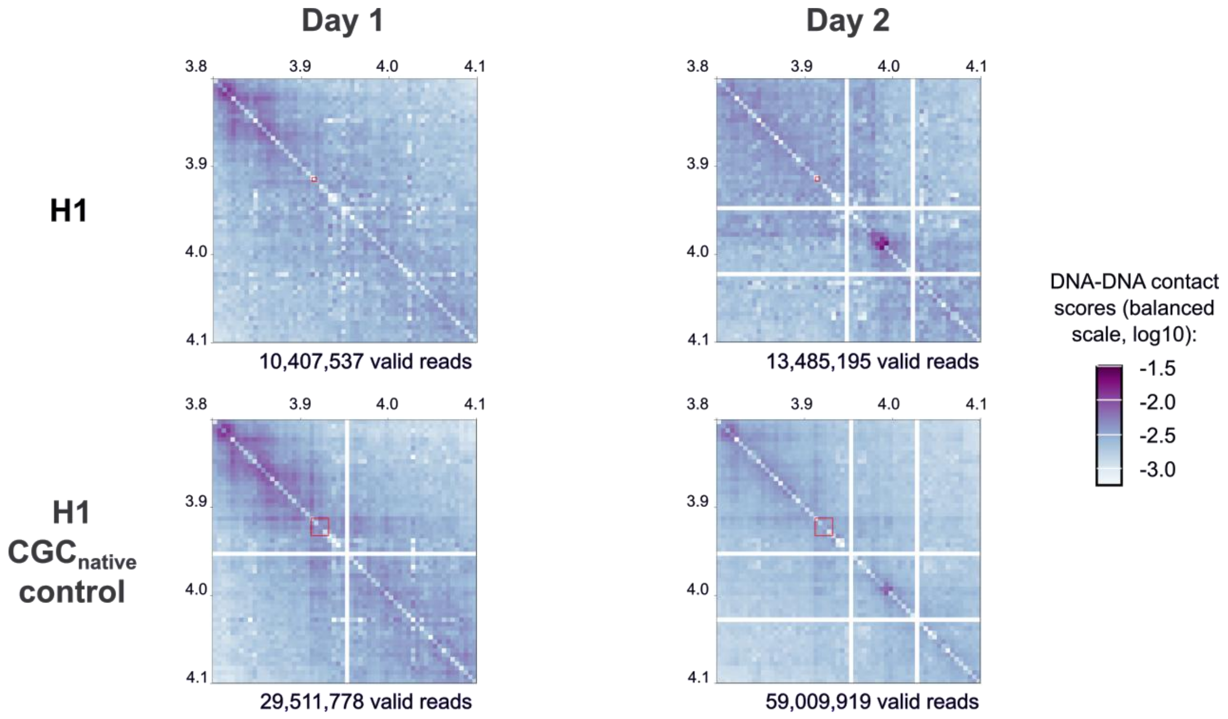


**Figure S4: Zoomed-in views of the *att*B-PhiC31 site in the parental strain without insertion and the H1-CGC$_{native}$-control strain**

Inset panels provide a zoomed-in view of a 300 kb region at 5 kb resolution, highlighting the *att*B-PhiC31 site, with genomic positions indicated in Mb. For the H1-CGC$_{native}$-control strain, the data represent merged results from 3 to 4 independent experiments shown in **Supplementary Figure S2**. The number of valid reads for each complete matrices, from which these zoomed-in regions are derived, is provided. The red squares mark largely the *att*B-PhiC31 site in the H1 strain and the pOSV806d vector in the H1-CGC$_{native}$-control strain, though they are visually enlarged for clarity. Introduction of the pOSV860Δ vector appears to induce a faint boundary, potentially linked to expression of the selection marker cassette. However, this subtle pattern does not develop into a full TID, unlike what is observed with the CGC cluster at Day 2 (**Fig.3**).

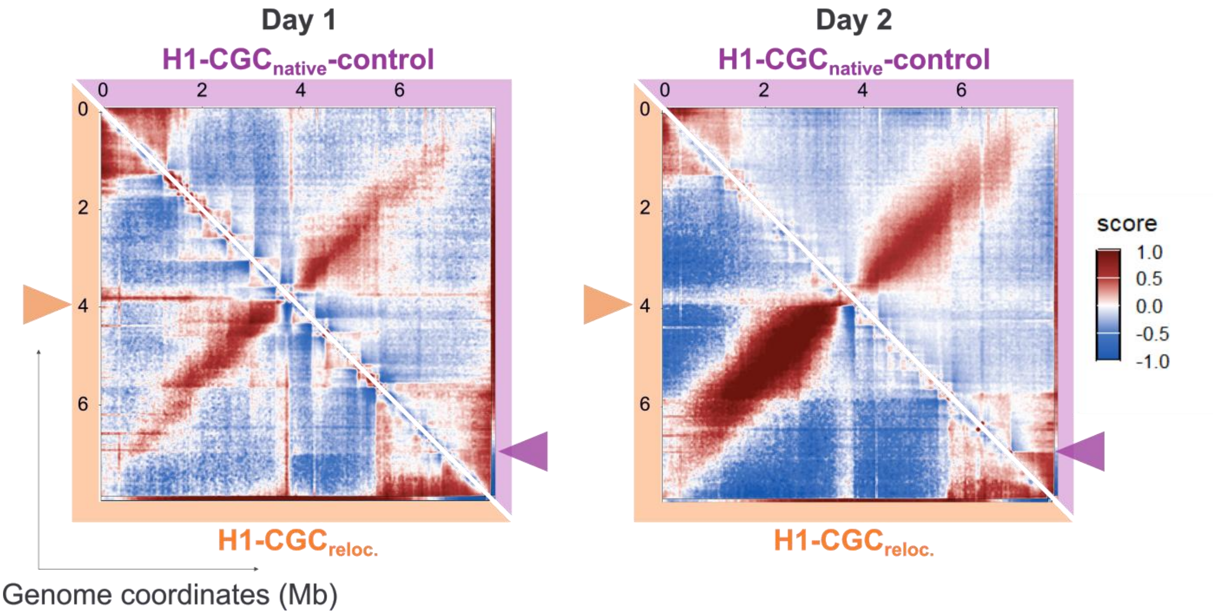


**Figure S5: Observed/Expected (OE) maps of the Hi-C matrices presented in Figure 3**
OE matrices are derived from balanced Hi-C matrices presented in **Figure 3** after detrending, using the OHCA pipeline (1). They visualize normalized chromatin interaction frequencies, showing how observed interactions deviate from expected values based on genomic distance. The color scale represents interaction strength: red indicates enriched interactions, white represents expected background levels, and blue shows depleted interactions. The triangles indicate the position of the CGC cluster in each strain.

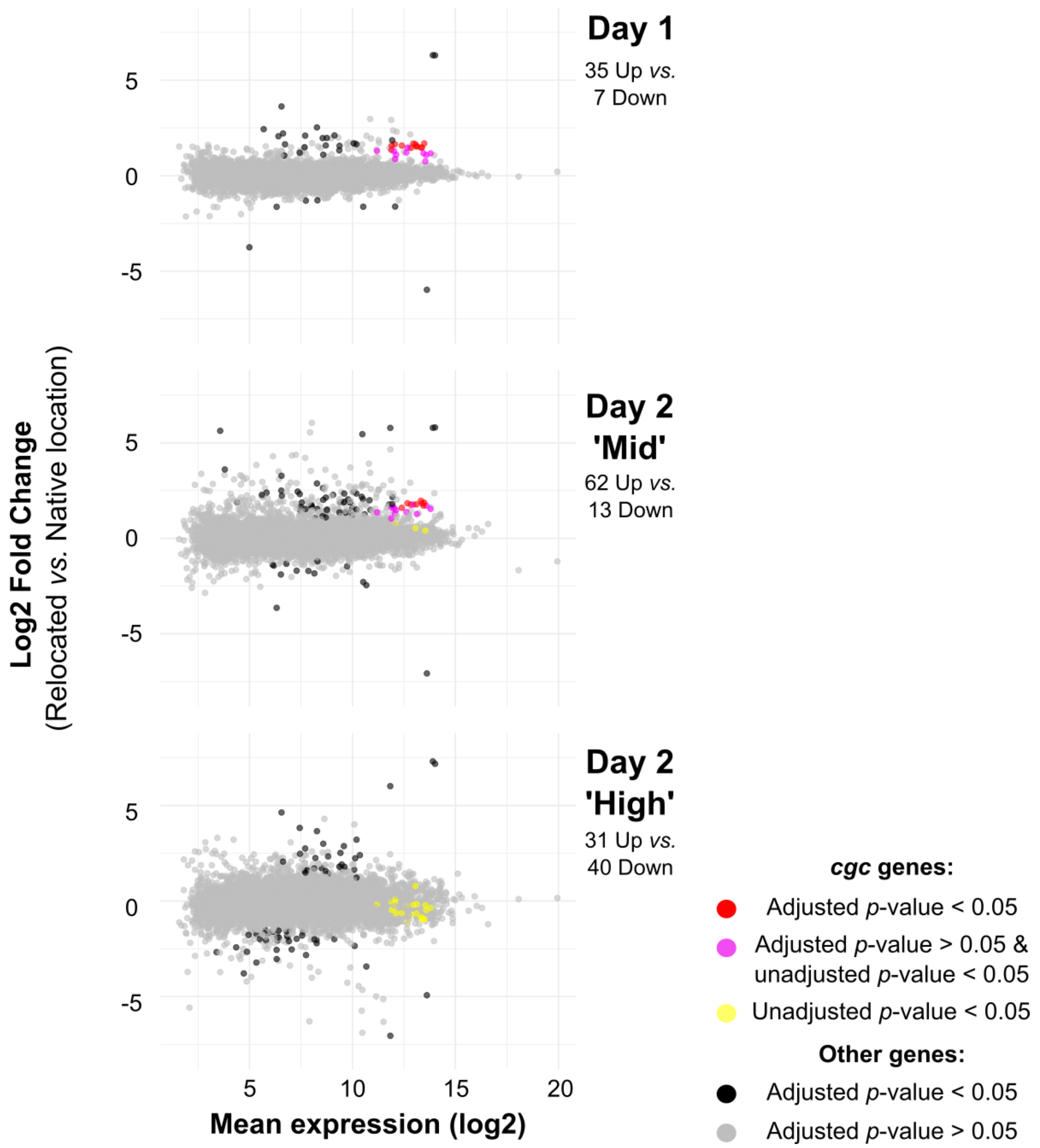


**Figure S6: Mean-average (MA) plot of differential gene expression in H1-CGC$_{reloc.}$ and H1-CGC$_{native}$ control strains over growth**

The results are presented for each gene, showing the average expression as a function of the fold change in the H1-CGC$_{reloc.}$ strain relative to the H1-CGC$_{native}$ control strain. The mean expression represents the average gene expression level across all replicates (combining both strains). The $\log_2$ value was calculated from this mean + 1 to avoid zero-related issues. Differentially expressed genes between the H1-CGC$_{reloc.}$ strain and the H1-CGC$_{native}$ control strain are highlighted for both the exponential phase (Day 1) and the transition toward the stationary phase at 'Mid' and 'High' stages (Day 2) in MP5 liquid medium, as indicated in the legend. Each panel shows the number of statistically over-expressed ('Up') or down-regulated ('Down') genes in the H1-CGC$_{reloc.}$ strain compared to the H1-CGC$_{native}$ control (adjusted *p*-value <0.05, EdgeR analysis). The full RNA-seq dataset is provided in **Supplementary Table S3**.

**Supplementary Table S1: Bacterial strains and plasmids used in this study**

| Name | Description | Reference |
|---|---|---|
| ***E. coli* strains** | | |
| DH5α | Host for general cloning and plasmid propagation | Promega |
| ET12567/pUZ8002 | Donor strain, whose conjugation helper non transmissible plasmid (pUZ8002) encodes kanamycin resistance, used for the conjugative transfer of DNA from *E. coli* to *Streptomyces* | (2) |
| ***Streptomyces ambofaciens* strains** | | |
| *S. ambofaciens* ATCC 23877 | RP3486 strain deposited at the ATCC by Rhône-Poulenc; (Sequenced genome: GCF_001267885.1_ASM126788v1) | (3), genome sequence (4), Pernodet-Lautru's lab collection |
| OSC2 | Derivative of *S. ambofaciens* ATCC 23877 devoid of pSAM2 | (5); Pernodet-Lautru's lab collection |
| H1 | Derivative of OSC2 with a scar-free deletion in the kinamycin (Δ*alpIABC*) | (6); Pernodet-Lautru's lab collection |
| CGCA018 | H1 strain with a deletion in the congocidine cluster *cgc22/cgc18*::*att3* (Δ*cgc22-cgc18*) | (7); Pernodet-Lautru's lab collection |
| H1-CGC$_{native}$-control | H1 strain harboring the pOSV806Δ plasmid integrated at the *att*B-PhiC31 site | This study |
| H1-CGC$_{reloc}$ | CGCA018 strain harboring the pCGC002 plasmid integrated at the *att*B-PhiC31 site | This study |
| H1-CGCΔ*cgc1*$_{reloc}$ | CGCA018 strain harboring the pCGC313 plasmid integrated at the *att*B-PhiC31 site | This study |
| ***Plasmids*** | | |
| pCGC002 | PhiC31 *att-int* integrative and conjugative BAC containing a 43.4 kb *S. ambofaciens* DNA fragment (7,187,197 to 7,230,572 bp position in *S. ambofaciens* ATCC23877 reference genome) including the complete *cgc* gene cluster and hygromycin resistance gene | (8) |
| pCGC313 | pCGC002-Δ*cgc1::att2* (Δ*cgc1* regulator gene) | (9) |
| pOSV806 | Conjugative plasmid carrying the hygromycin resistance gene and PhiC31 *att-int* system (Addgene 126606) | (10) |
| pOSV806Δ | pOSV806 vector lacking the cloning module (*amilCP* cassette) following NotI digestion and ligation | This study |

**Supplementary Table S2: Fold change in *cgc* gene expression during exponential phase (Day 1) in the H1-$CGC_{reloc.}$ and H1-$CGC_{native}$-control strains, before and after normalization to MFA counts**

| gene | Before normalization | After normalization* |
|---|---|---|
| *cgc 22* | 2.9 | 1.1 |
| *cgc21* | 2.9 | 1.3 |
| *cgc20* | 2.8 | 1.3 |
| *cgc1* | 1.8 | 0.9 |
| *cgc2* | 2.4 | 1.2 |
| *cgc3* | 3.0 | 1.4 |
| *cgc4* | 3.2 | 1.5 |
| *cgc5* | 2.7 | 1.3 |
| *cgc6* | 2.7 | 1.2 |
| *cgc7* | 3.6 | 1.6 |
| *cgc8* | 2.4 | 1.2 |
| *cgc9* | 2.8 | 1.5 |
| *cgc10* | 2.6 | 1.3 |
| *cgc11* | 2.5 | 1.1 |
| *cgc12* | 2.1 | 1.0 |
| *cgc13* | 2.8 | 1.2 |
| *cgc14* | 1.8 | 0.9 |
| *cgc15* | 2.9 | 1.4 |
| *cgc16* | 2.2 | 1.0 |
| *cgc17* | 2.0 | 1.1 |
| *cgc18* | 2.8 | 1.0 |
| *cgc19* | 2.9 | 0.9 |
| **mean** | **2.6** | **1.2** |
| **SEM** | **0.1** | **0.04** |

RNA-seq counts for each gene were normalized to the corresponding MFA counts prior to EdgeR analysis to determine fold changes between conditions. The analysis was performed using four independent biological replicates, with two strains analyzed per replicate.
